# Pharmaceutical TAS2R14 Agonists Display Diverse Potency, Efficacy, and Binding-Site Sensitivity

**DOI:** 10.64898/2026.06.17.732860

**Authors:** Shir Eyal, Nitsan Dallal, Alon Rainish, Evgenii Ziaikin, Einav Malach, Masha Y Niv

## Abstract

Bitter taste receptors (TAS2Rs) are G-protein coupled receptors that detect chemically diverse compounds, including many clinically used drugs. TAS2R14 is expressed in many extraoral tissues and is activated by hundreds of ligands, including pharmaceutical drugs. Recent cryo-EM structures revealed a previously unrecognized intracellular binding pocket in TAS2R14, raising new questions regarding ligand binding modes.

Here, we investigated the activation of TAS2R14 by Tamoxifen, Carbimazole, and Lidocaine using cell-based assays measuring proximal G-protein recruitment (BRET2) and downstream signaling (IP-One). Tamoxifen and Carbimazole activated TAS2R14 with EC50 values in the low micromolar range, whereas Lidocaine required substantially higher concentrations. Targeted receptor mutations were used to evaluate the contribution of extracellular and intracellular binding regions to agonist activity. Carbimazole and Lidocaine showed greater dependence on the intracellular and extracellular positions, respectively, while Tamoxifen displayed assay-dependent, but overall modest sensitivity to the tested mutations. Thus, although existing drugs can activate TAS2R14 through distinct binding modes, TAS2R14-directed repurposing will depend on whether effective local receptor concentrations can be achieved through appropriate delivery strategies.

## 1) Introduction

G-protein coupled receptors (GPCRs) represent the largest family of membrane receptors in mammals, playing crucial roles in numerous physiological processes. These receptors are activated by diverse ligands, and the resulting conformational changes trigger intracellular signaling cascades^1^. GPCRs’ widespread involvement in vital biological functions makes them a major target in drug development, with around 36% of FDA-approved drugs designed to modulate their activity^2,3^. In the taste system, GPCRs are responsible for signal transduction in sweet, umami, and bitter taste pathways through specific G protein complexes^1^.

Bitter taste receptors (TAS2Rs), a subfamily of GPCRs, enable the detection of a wide variety of bitter compounds^4^, often as a means to avoid ingestion of potentially harmful substances^1^. While bitterness has long been associated with toxicity detection, this interpretation is evolving, as some bitter compounds like flavonoids have beneficial properties, indicating that bitterness is a weak indicator of toxicity^5–7^.

The bitter taste of many clinical drugs presents a major problem for patient compliance, especially in children and the elderly. Since bitterness is often evaluated only at late stages of drug development, the identification of intensely bitter compounds can result in expensive reformulations, increased animal testing, and significant delays in market approvals^8,9^. While pharmaceutical technologies are commonly applied to mask these unpleasant tastes^10^, bitterness should not be considered only in terms of formulation. The expression of TAS2Rs in extraoral tissues, including the lungs, gut, and brain, suggests that bitter drugs may exert physiologically relevant effects beyond taste perception^11–14^.

TAS2R14 is the most extensively characterized bitter taste receptor and is notable for its ability to recognize a wide range of structurally diverse ligands, including both natural and synthetic compounds, many of which are pharmacologically active drugs^15^. According to BitterDB^4^, the receptor currently has 385 known agonists. This broad ligand spectrum is especially intriguing given TAS2R14 expression outside the oral cavity^11–14^, where its activation may influence non-gustatory physiological processes.

Here, we focus on pharmaceutical drugs previously implicated in TAS2R14 activation^16,17^. Many of the agonists listed in BitterDB^4^ originate from large-scale screening studies that evaluated receptor activation using calcium mobilization assays^15,16,18^. These efforts substantially expanded the known TAS2R14 ligand repertoire, but potency and efficacy, was measured for only part of the ligands.

We investigated the concentration-dependent activation of TAS2R14 by representative pharmaceutical agonists. We hypothesize that pharmaceutical TAS2R14 agonists differ substantially in their potency, efficacy, and dependence on specific receptor interactions. Determining these properties may help evaluate the physiological relevance of TAS2R14 activation by clinically used drugs, the possibility that some of their activity is mediated via TAS2R14^20^ and evaluating their potential for drug repurposing^18,19^.

Cryo-EM structures of agonist-bound TAS2R14 revealed a novel intracellular binding pocket^21–24^ that appears to serve as the primary site of interaction for several agonists. This discovery raised new questions regarding the functional contribution of this intracellular binding site relative to the extracellular binding site, and whether different ligands preferentially engage one or both regions of the receptor. We recently showed that TAS2R14 antagonists differ in their dependence on the extracellular and intracellular pockets for receptor inhibition^25^.

To choose candidates representing these different options, we used docking of known agonists to the extracellular and measures the downstream output intracellular pockets in the cryo-EM structure of TAS2R14 (PDB: 8RQL^21^).

To capture both downstream signaling and proximal G-protein engagement, receptor activity was evaluated using BRET2 assays based on the TRUPATH platform, which directly monitors G-protein coupling, and the IP-One assay, which measures downstream IP1 accumulation.^25–27^

## 2) Materials and Methods

### Cell culture

HEK293T cells were used for all experiments. Cells were maintained in Dulbecco’s Modified Eagle Medium (DMEM) supplemented with 10% fetal bovine serum (FBS), 1% L-glutamine, and 0.2% penicillin-streptomycin. Cultures were incubated at 37 °C in a humidified atmosphere containing 5% CO₂. Cells were passaged every 3-4 days upon reaching approximately 80% confluence.

### Compounds

Materials were purchased from the following suppliers: Thermo Fisher Scientific (USA; Lidocaine HCl, Carbimazole), Sigma (USA; Flufenamic acid), Aaron Chemicals (USA; Tamoxifen), Gibco (USA; DMEM, FBS), Revvity (France; IP-One HTRF kit), BLDpharm (China), and Mirus Bio (USA; TransIT-293 reagent). Some of the tested compounds were initially dissolved in DMSO (with final concentration not exceeding 1%) and others in each assay’s buffer. On the day of each experiment, serial dilutions were prepared in the relevant buffer for each assay.

### IP-One assay

To assess TAS2R14 activity, HEK293T cells were transiently transfected at 50–80% confluence using Mirus TransIT-293 (Mirus Bio, USA) in a 1:3 DNA-to-reagent ratio. Each transfection included 2 µg of a pcDNA3.1 plasmid encoding a modified version of wild-type (WT) TAS2R14 or mutated TAS2R14 (with a FLAG tag, and the first 45 residues of the rat somatostatin receptor 3 to enhance membrane trafficking), along with 1 µg of a plasmid expressing the chimeric Gαqi5 protein (Addgene plasmid #24501, RRID: Addgene_24501).

24h post-transfection, cells were resuspended in 10% DMEM and plated in 384-well microplates (Greiner). After allowing cell adherence for an additional 24 h, the medium was replaced with stimulation buffer (provided with the Revvity IP-One Gq kit), which includes 50 mM LiCl to inhibit IP1 degradation.

The cells were exposed to the tested compounds and incubated for 150 minutes to promote IP1 accumulation.

To terminate signaling and initiate detection, cells were lysed with detection reagents (IP1-d2 and anti-IP1 cryptate-TB conjugates in kit lysis buffer). Fluorescence was measured using a FRET-based approach, with emission at 665 nm (acceptor) and 620 nm (donor). FRET ratios were normalized to a baseline (0%) and the maximum agonist response (100%). Dose-response curves were fitted using a four-parameter logistic model in GraphPad Prism (GraphPad Software, La Jolla, CA) to determine ligand potency (EC₅₀) and efficacy (Emax, relative to each agonist). Each compound was tested in 3 independent experiments, with concentrations evaluated in triplicate.

### BRET2 assay

BRET2 assays were performed using TRUPATH resources^26^. HEK293T cells were transiently transfected at 50–80% confluence using Lipofectamine 3000 (Thermo Fisher Scientific) according to the manufacturer’s protocol. Each transfection included 1 µg of a pcDNA3.1 plasmid encoding either WT or mutated TAS2R14 (with an N-terminal HA signal peptide, FLAG tag, and the first 45 residues of the rat somatostatin receptor 3), along with plasmids encoding TRUPATH biosensor components: Gα-gustducin fused to Renilla luciferase 8 (RLuc8), Gβ1, and Gγ13 fused to GFP2 at a 1:1:1:1 ratio.

Plates were incubated for 24 h at 37 °C and 5% CO_2_. Transfected cells were harvested and seeded at 50,000 cells/well into poly-D-lysine treated 96-well flat bottom white culture plates and grown for 24 h at 37 °C and 5% CO_2_. After allowing adherence for an additional 24 h, Cells were washed and then incubated in 80 µL assay buffer (1x HBSS, 24 mM HEPES, 3.96 mM NaHCO3, 1.3 mM CaCl2, 1 mM MgSO4 and 10% w:v BSA) for 1 h. Following incubation, coelenterazine 400a (CTZ400a, NanoLight) was added to each well to a final concentration of 10 µM, and plates were equilibrated briefly at 37 °C. Immediately after CTZ400a addition, BRET² measurements were initiated and recorded continuously for 5 min to establish a baseline signal. Following baseline acquisition, 10 µL of ligand solution or vehicle was added to each well while BRET recording continued. For stimulation experiments, the added solution contained the tested compound at the indicated concentrations. BRET signals were monitored for several minutes after ligand addition to capture receptor-dependent activation of the G protein complex.

Measurements were performed using a Clariostar® Plus plate reader (BMG LabTech, Germany) equipped with emission filters centered at 515–30 nm for GFP2 and 410–80 nm for RLuc8. BRET ratios were calculated as the emission signal of GFP2 divided by the emission of RLuc8. ΔBRET values were obtained by correcting raw BRET signals using vehicle-treated wells.

Responses were normalized to vehicle-treated wells (0%) and to the maximal response induced by a reference agonist (100%). Dose–response curves were fitted using a four-parameter logistic model in GraphPad Prism (GraphPad Software, La Jolla, CA) to determine ligand potency (EC₅₀) and efficacy (Emax). Each compound was tested in at least three independent experiments, with concentrations evaluated in duplicate or triplicate wells.

### Docking of known ligands

To evaluate the binding-site preference of TAS2R14 ligands, molecular docking was performed against both the extracellular and intracellular binding pockets of the solved receptor structure. The receptor (PDB: 8RQL^21^) was prepared with the protein preparation wizard (Schrödinger Release 2025-1: Protein Preparation Workflow; Epik, Schrödinger, LLC, New York, NY, 2024; Impact, Schrödinger, LLC, New York, NY; Prime, Schrödinger, LLC, New York, NY, 2025.) available within the Maestro Schrödinger Suite (Schrödinger Release 2025-1: Maestro LLC, New York, NY, 2025). The TAS2R14’s ligands known while performing this step were collected from BitterDB^27^ and prepared with the LigPrep tool (Schrödinger Release 2025-1: LigPrep, Schrödinger, LLC, New York, NY, 2025). The grids were generated based on the solved structure ligand locations (extracellular and intracellular FFA). All the ligands were docked independently into both sites using the OPLS3e force field^28^ with the Glide XP protocol^29^ available within Schrödinger (Schrödinger Release 2025-1: Glide XP protocol, Schrödinger, LLC, New York, NY, 2025). Docking parameters were kept identical for both grids to enable direct comparison of docking scores. Docking scores obtained from Glide XP were used to assign ligand binding-site preferences using predefined absolute and relative score criteria. Ligands were classified to binding site based on docking score thresholds and site-specific score differences, with FFA used as a reference dual-site ligand.

## 3) Results

### 3.1) Computational prediction of ligand binding site preferences

We docked 385 known TAS2R14 ligands (collected from BitterDB^4^) to the top and the bottom binding site of the receptor independently, starting from the 8RQL.pdb structure^21^ using Glide XP (Maestro, Schrödinger)^30^. Docking scores were used to classify ligands according to their predicted binding site using two complementary criteria. First, an absolute docking score threshold was used to exclude weak or failed docking poses. Docking scores higher than −5.0 were considered indicative of poor binding; ligands that failed to reach this threshold at a given site were classified as non-binders for that site.

Second, relative site preference was determined using a score-difference (Δ score) criterion. FFA, which has been experimentally observed in the intracellular binding site of some TAS2R14 cryo-EM structures^22^, while binding to the extracellular site alongside the intracellular site has been reported in only one structure^21^, was used as a reference compound. Because experimental structures indicate that FFA is capable of occupying both binding pockets under at least some conditions, and given that FFA has more frequently been observed in an intracellular-only binding mode, a conservative threshold of Δ ≤ 2.0 (Δ = 2.3 for flufenamic acid) was applied for predicted ligand “dual-site” binder: Ligands that achieved favorable docking scores (≤ −5.0) at both sites and exhibited an absolute Δ score below 2.0 were classified as potential dual-site binders. Ligands with favorable scores at both sites but a Δ score greater than 2.0 were assigned to the site with the stronger (more favorable) docking score. If only one site yielded a favorable docking score, the ligand was classified as site-selective for that pocket with no delta consideration. Ligands that failed to reach the docking threshold at both sites were categorized as weak binders and classified as “other” (Fig. 1a). This classification scheme enabled systematic assignment of binding-site preferences across the full ligand dataset and facilitated a predicted comparative analysis of extracellular versus intracellular binding tendencies within the TAS2R14 ligand repertoire.

**Figure 1.**
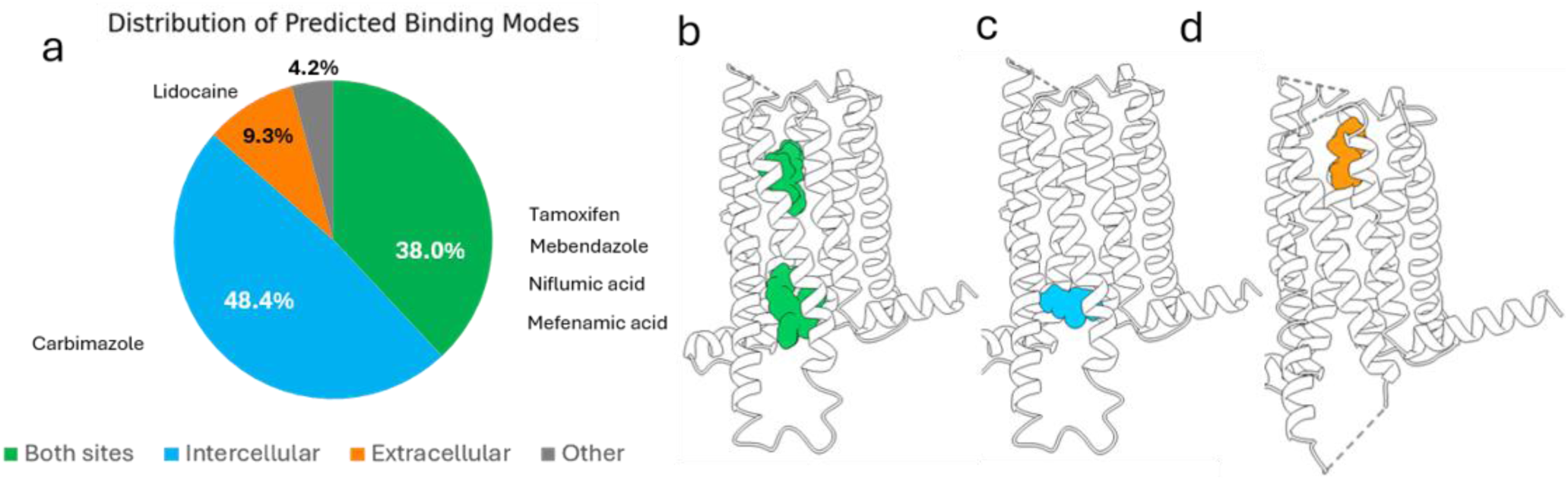
(a) Distribution of predicted ligand binding sites based on docking analysis **-** Pie chart showing the proportion of docking poses predicted to bind to both extracellular and intracellular sites (green), the intracellular site only (blue), the extracellular site only (orange), and “other” (grey). Selected compounds are marked as they exhibit distinct site preferences. (b-d) docking poses of (b), Tamoxifen (c), Carbimazole (d), and Lidocaine.

Following the classification of ligands, 6 representative compounds from the different categories were selected for experimental investigation (Fig. 1b-d). All compounds were first tested in MOCK-transfected HEK293T cells using the IP-One assay to exclude non-specific effects (Supplementary Fig. S1). Three compounds, Carbimazole^31^, Tamoxifen^32^ and Lidocaine^17^ showed no detectable activity in MOCK-transfected cells and were selected for further characterization. Tamoxifen is a non-steroidal anti-oestrogen, which was originally developed as a potential contraceptive and later became the first targeted therapy for breast cancer due to its ability to target the oestrogen receptor^32^, was predicted to bind both extracellular and intracellular pockets of TAS2R14; Carbimazole, an antithyroid drug and a prodrug of methimazole, which inhibits thyroid hormone synthesis by blocking the iodination of tyrosine residues in thyroglobulin, a process mediated by thyroid peroxidase^31^, was predicted to bind to the intracellular pocket of TAS2R14, and Lidocaine, a widely used local anesthetic that blocks voltage-gated sodium channels to inhibit pain signaling^17^, was predicted to bind to the extracellular pocket of TAS2R14.

### 3.2) Functional characterization of selected TAS2R14 agonists

Next, we examined how the selected drug agonists functionally activate TAS2R14 and how mutations in the two binding pockets affect receptor signaling.

As a reference compound we used FFA, a nonsteroidal anti-inflammatory drug (NSAID) that inhibits cyclooxygenase-2 (COX2) and is one of the most potent known agonists of TAS2R14^18^, for which cryo-EM structure of its complex with TAS2R14 has been resolved in our laboratory and others^21,22^.

Receptor activation was quantified using two experimental approaches: BRET2 and IP-One assays, which capture different stages of GPCR signaling. The BRET2 assay provides a readout of proximal G-protein engagement by detecting early events directly at the receptor-transducer interface^26^, whereas the IP-One assay quantifies downstream Gq-pathway activation by measuring the cellular accumulation of IP1^33,34^.

Dose-response curves were then obtained for Carbimazole, Tamoxifen, and Lidocaine in WT TAS2R14 using both IP-One and BRET2 assays, normalized to the maximal activation by the reference agonist FFA. All three compounds activated TAS2R14, with Tamoxifen producing the strongest response, whereas Carbimazole and Lidocaine displayed lower efficacy compared to FFA (Fig. 2). Lidocaine was the least potent in the BRET2 assay and did not have a reproducible effect in IP-One assay.

**Figure 2:**
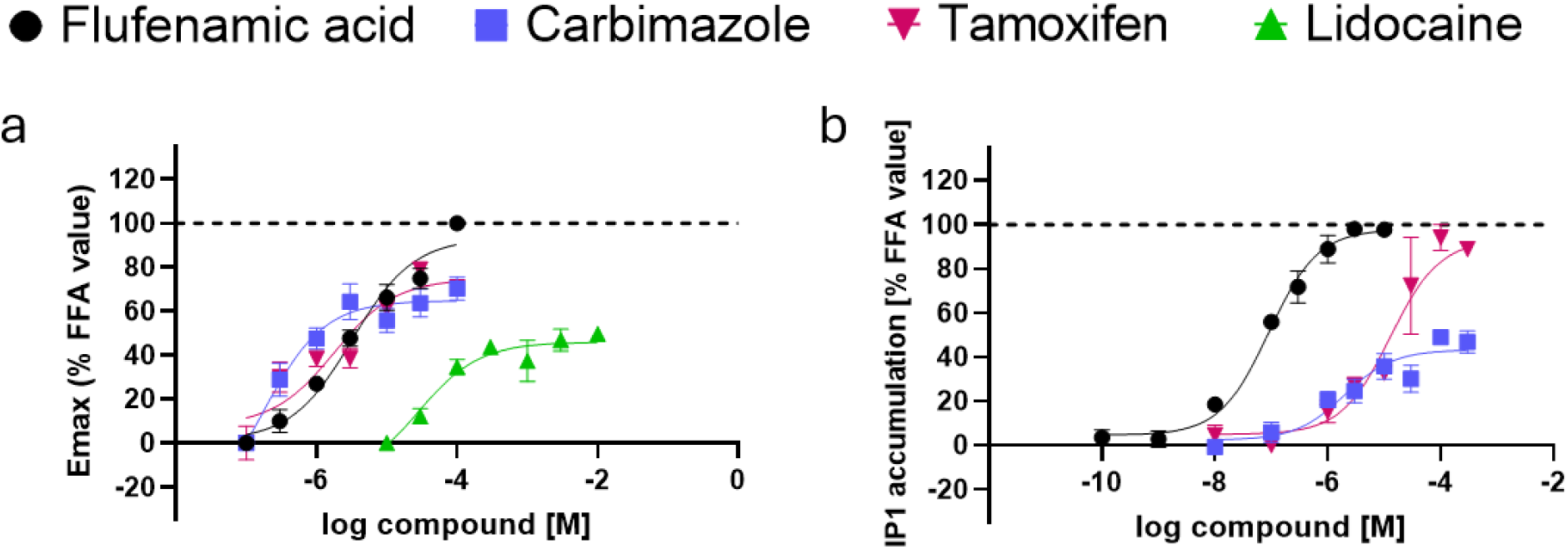
Activation of WT TAS2R14 by selected agonists in HEK293T cells, values represent the mean ±SEM. 3 biological repeats. FFA in black, Carbimazole in purple, Tamoxifen in pink and Lidocaine in green (A) BRET assay results-Measured Δ BRET (RLuc8/Gβ1γ2-GFP2) in WT. (B) IP-One results-measured Gαqi5 mediated accumulation of IP1% of FFA in WT.

To investigated the relative contributions of the extracellular and intracellular binding sites, we examined receptor activation following mutations at two key residues previously implicated in ligand interactions^21–24^: S265A^7.38^ in the extracellular binding site and S194A^5.54^ in the intracellular binding site (Fig. 3).

**Figure 3:**
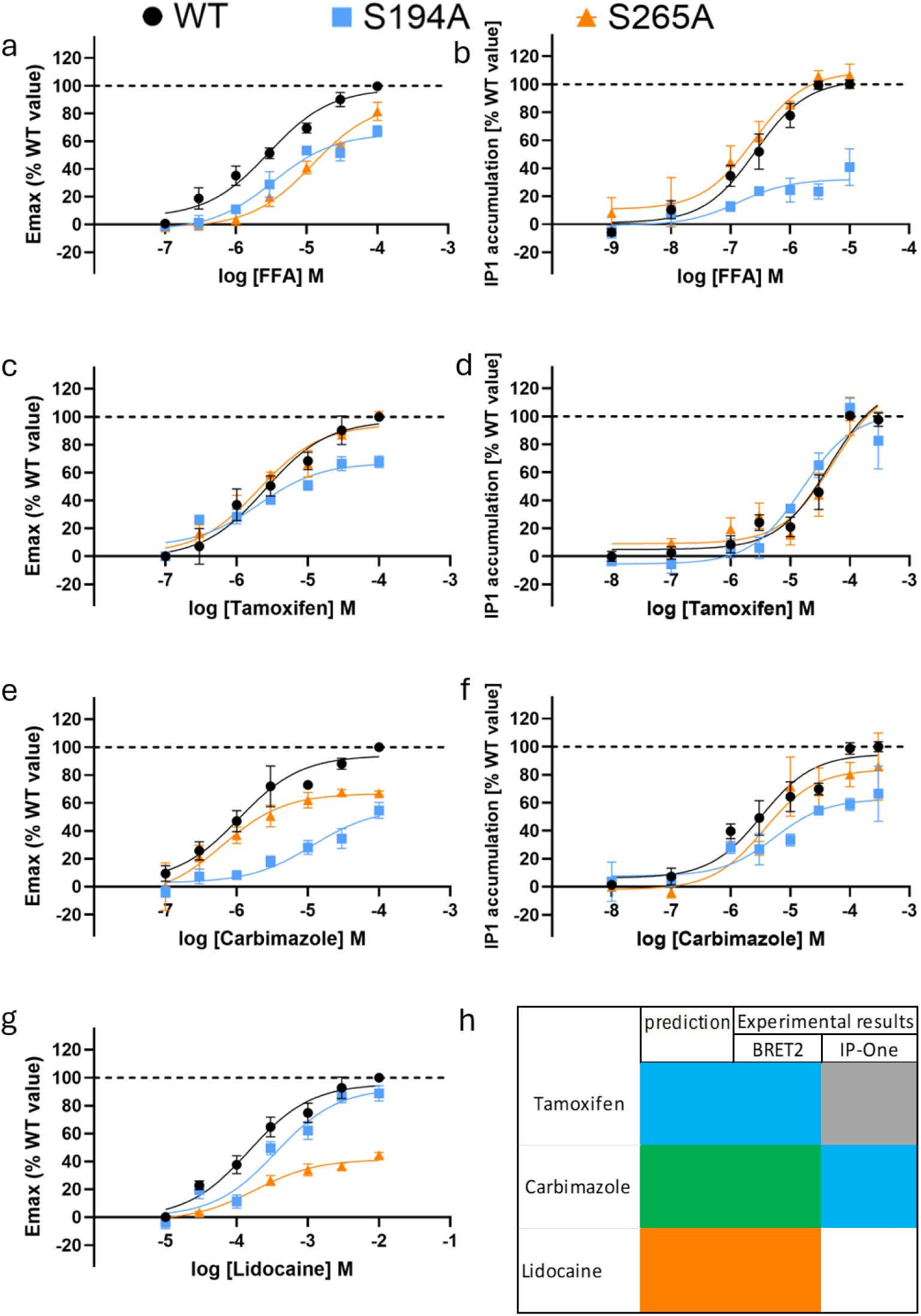
Activation of S194A and S265A mutants. HEK293T cells transfected with WT or mutated TAS2R14. Values represent the mean ±SEM. 3 biological repeats. WT in black, S265A^7.38^ in orange and S194A^5.54^ in blue. (a,c,e,g) BRET2 assay results (b,d,f) IP-One results-measured Gαqi5 mediated accumulation of IP1% of each agonist in WT (a,b) FFA (c,d) Tamoxifen (e,f) Carbimazole (g) Lidocaine. (h) Summary of predictions and experimental results, Blue indicates intracellular interaction, orange extracellular interaction, green both sites, and gray no site-specific effect. All EC_50_ and E_max_ values are presented in table 1.

**Table 1:**
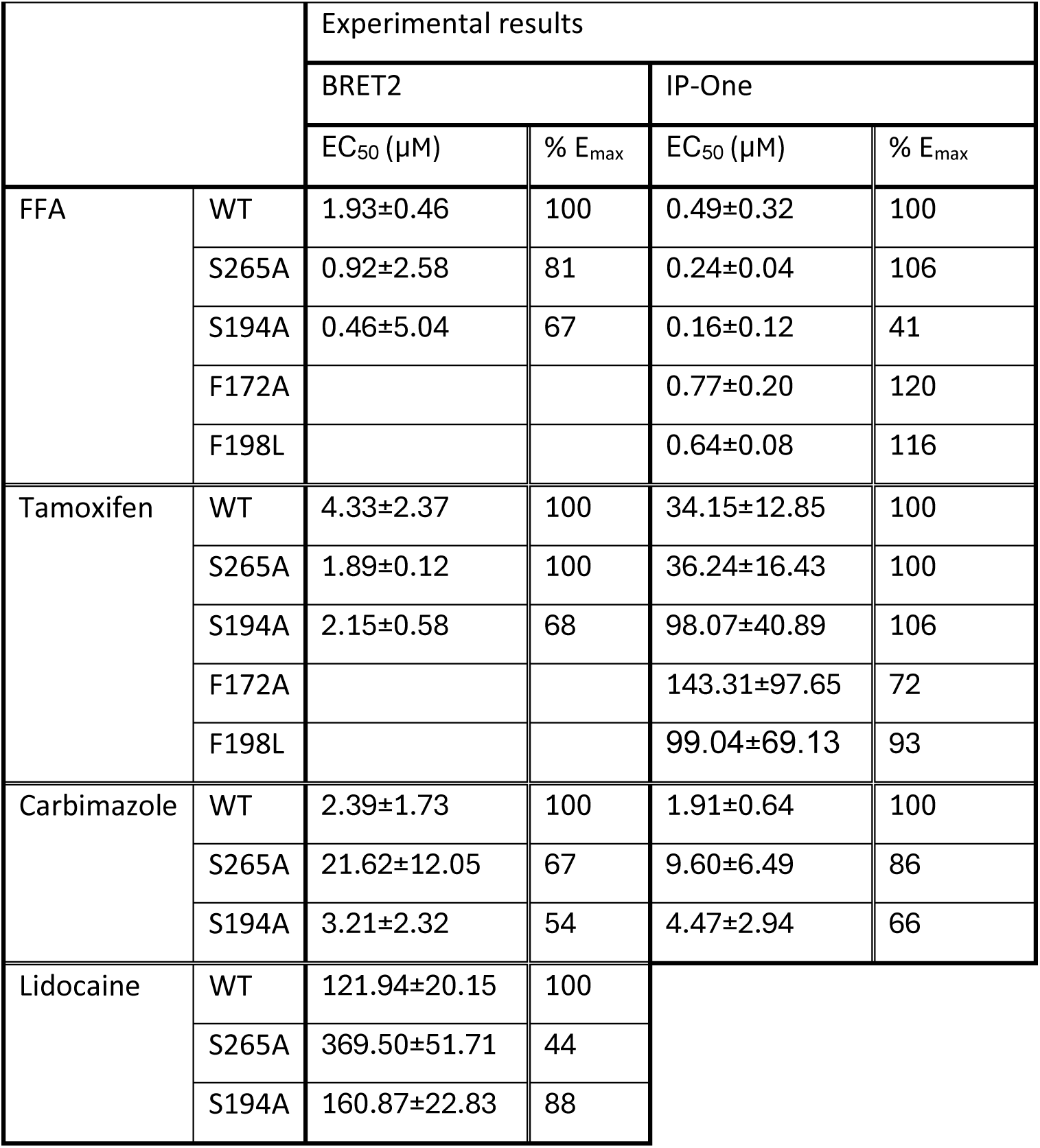
Potency (EC₅₀) and efficacy (E_max_) of the tested compounds at WT and mutant TAS2R14 receptors.

For FFA, consistent with previous studies, mutation in the intracellular binding site (S194A^5.54^) substantially reduced receptor activation in both the BRET2 and IP-One assays (E_max_= 67%, 41%, respectively, table 1). In contrast, mutation in the extracellular binding site (S265A^7.38^) had only a modest effect on maximal receptor activation, observed only in the BRET2 assay (E_max_= 81%) (Fig. 3a,b). The importance of the intracellular site is consistent with cryo-EM structures showing FFA binding in the intracellular pocket^21,22^. While FFA has also been observed in the extracellular pocket in one reported structure^21^, previous mutagenesis studies similarly identified additional intracellular-pocket residues contributing to FFA-mediated receptor activation.

For Carbimazole, in the BRET2 assay, both mutations reduced G-protein recruitment, with a stronger decrease observed for S194A^5.54^ than S265A^7.38^ (E_max_=54% and 67%, respectively). In the IP-One assay, S265A^7.38^ showed minimal difference from WT activation (E_max_=86%), whereas S194A^5.54^ resulted in a more substantial reduction in receptor activation (E_max_= 66%) (Fig. 3c,d). Despite differences between the assay readouts, both approaches revealed that disruption of the intracellular binding pocket had a greater impact on carbimazole-mediated activation than disruption of the extracellular site, aligning with the computational prediction.

For Lidocaine, in the BRET2 assay, the intracellular mutation had a minimal effect on TAS2R14 activation (E_max_= 88%), whereas the extracellular mutation caused a notable decrease in activation (E_max_= 44%) (Fig.3 g). These findings indicate that Lidocaine-mediated activation is more sensitive to alterations in the extracellular binding site, consistent with the docking prediction. While this compound produced measurable responses in the BRET2 assay, no detectable signal was observed in the IP-One assay. This discrepancy may reflect several non-mutually exclusive factors, including the high concentrations required to elicit a response (up to 10 mM), which could introduce non-specific or cytotoxic effects, as well as inherently weak coupling of Lidocaine to downstream Gq-mediated signaling.

In contrast to Lidocaine, docking predicted that Tamoxifen could interact with both binding sites. Experimentally, the two assays produced different outcomes and, notably, Tamoxifen was the only ligand for which the EC_50_ values differed substantially between assays, with BRET2-derived EC_50_ values being approximately one order of magnitude lower than those obtained in the IP-One assay. In the BRET2 assay, S194A^5.54^ resulted in a moderate reduction in G-protein recruitment (E_max_ = 68%), whereas S265A^7.38^ produced a response comparable to WT (Fig. 3e,f). In the IP-One assay, neither mutation significantly altered Tamoxifen-induced receptor activation relative to WT. This divergence between the two assays may suggest that intracellular interactions contribute to proximal G-protein coupling, whereas downstream signaling remains largely intact despite disruption of either binding site.

Given the limited effect of S194A^5.54^ and S265A^7.38^ on Tamoxifen-induced activation, we selected two additional residues that appeared in the docking-derived interaction maps as Tamoxifen-interacting positions (Fig. S2): F172^4.62^ is in the extracellular binding pocket, and F198^5.58^ lies in the intracellular binding pocket. These residues were evaluated in the IP-One assay, with FFA tested in parallel as a reference agonist (Fig. 4a,b).

**Figure 4:**
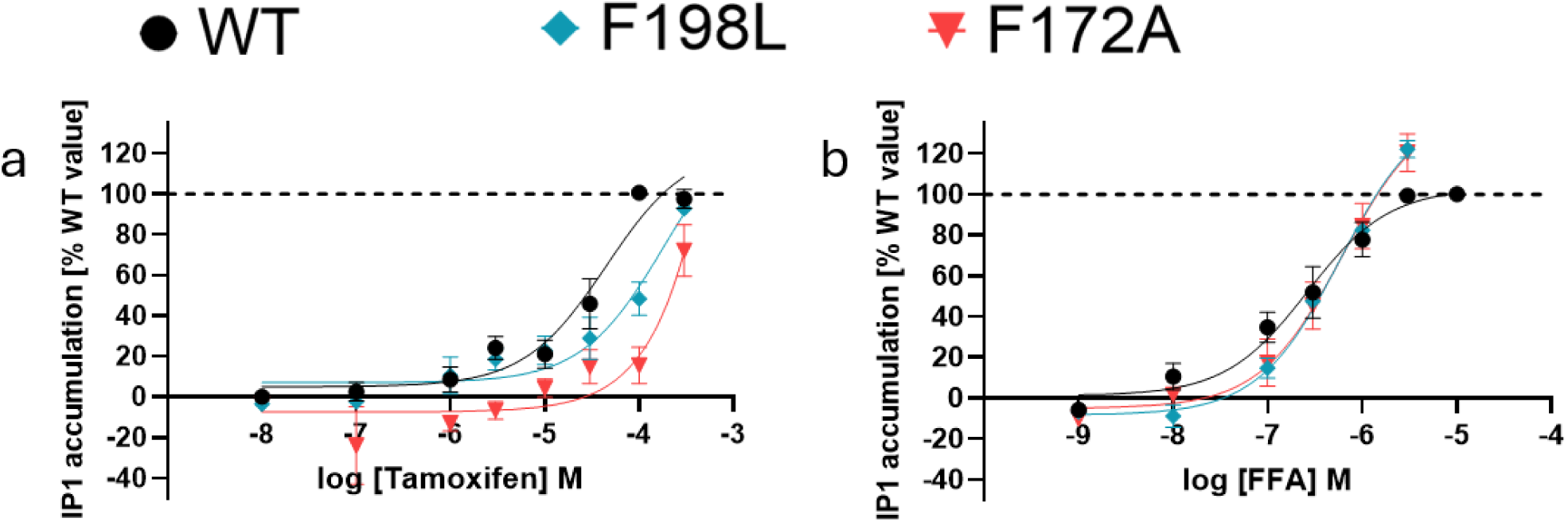
Effects of F172A and F198L mutants by Tamoxifen and FFA-. HEK293T cells transfected with WT or mutated TAS2R14, measured Gαqi5 mediated accumulation of IP1% of each agonist in WT Values represent the mean ±SEM. 3 biological repeats. WT in black, F172A^4.62^ in red and F198L^5.58^ in turquoise. (a) Tamoxifen (b) FFA. All EC50 and Emax values are presented in table 1.

F172A^4.62^ reached 72% of the WT maximal response, with an EC_50_ of 143.3 ± 97.7μM, compared with 34.1 ± 12.9 μM for WT. F198L^5.58^ showed near WT-like maximal activation, reaching 93% of the WT response, with an EC_50_ of 99.0 ± 69.1 μM. Thus, both mutations shifted the response to higher concentrations, while substantial receptor activation was retained.

FFA activity was largely preserved in both mutants, indicating that the effects observed for Tamoxifen are not a general consequence of impaired receptor function.

The IP-One results suggest that F172^4.62^ and F198^5.58^ contribute to Tamoxifen recognition, with F172 playing a more important role. The preserved activation in mutations affecting both the extracellular and intracellular pockets is consistent with a binding mode in which Tamoxifen is supported by several contacts across the pocket environment, rather than by a single dominant anchor. In this view, the EC_50_ shifts may reflect weaker ligand binding, whereas the retained E_max_ indicates that the receptor can still reach a signaling-competent active state after local perturbation of the predicted interaction network.

## 4) Discussion

In this study, we quantitatively characterized three pharmaceutical TAS2R14 agonists, Tamoxifen, Carbimazole, and Lidocaine. While these compounds had previously been reported to activate TAS2R14, information regarding their potency and efficacy was limited. Using both BRET2 and IP-One assays, we determined their concentration-response relationships and examined how mutations in the extracellular and intracellular binding pockets influence receptor activation. The combined functional and computational analyses revealed substantial differences in both signaling behavior and binding-site dependence among the tested agonists.

### 4.1) BRET2 and IP-One capture distinct stages of TAS2R14 signaling

Although the overall trends observed in the BRET2 and IP-One assays were generally consistent, several ligand-mutation combinations produced assay-dependent effects. These differences likely reflect the distinct stages of GPCR signaling measured by each approach. The BRET2 assay monitors proximal G-protein activation by detecting ligand-induced rearrangement or dissociation of G protein subunits^26^, whereas the IP-One assay reflects the downstream accumulation of IP1 generated following phospholipase C activation in the Gq signaling pathway^33,34^.

Accordingly, alterations in receptor G-protein coupling may be more readily detected by BRET2, whereas downstream responses may remain partially preserved due to signal amplification and temporal integration of signaling outputs. In addition, the two assays differ in the G proteins they engage, with BRET2 reporting coupling to Gα-gustducin and IP-One reflecting signaling mediated by the chimeric Gαqi5. Differences between the assays may therefore also arise from ligand-specific G protein coupling preferences. Consistent with this, previous studies have shown that BRET-based assays can detect ligands as partial agonists, whereas downstream readouts such as cAMP accumulation may present the same ligands as full agonists^35^.

Tamoxifen provided the clearest example of this phenomenon. Mutation of the intracellular binding site reduced receptor activation in the BRET2 assay, while having little effect on the IP-One response. Thus, Tamoxifen appeared more potent in the BRET2 assay than in the IP-One assay. These observations may indicate that the consequences of disrupting ligand-receptor interactions can differ depending on the stage of the signaling pathway being measured. One possible explanation is that partial reductions in coupling efficiency can be detected in BRET2, whereas downstream signaling remains relatively preserved.

### 4.2 Pharmaceutical agonists differ in potency and binding-site dependence

Tamoxifen, Carbimazole, and Lidocaine have previously been reported as TAS2R14 agonists, yet quantitative pharmacological characterization of these compounds has remained limited. To our knowledge, this study provides the first systematic comparison of their concentration-response relationships using both proximal G-protein recruitment and downstream signaling assays. The EC_50_ and E_max_ values determined here therefore provide a quantitative framework for comparing the activity of pharmaceutical TAS2R14 agonists across distinct stages of receptor signaling.

The tested compounds differed substantially in both potency and sensitivity to receptor mutations. Tamoxifen and Carbimazole activated TAS2R14 at low micromolar concentrations, whereas Lidocaine required considerably higher concentrations to elicit receptor activation.

The mutagenesis experiments revealed distinct patterns of binding-site dependence among the tested agonists. Carbimazole was more sensitive to disruption of the intracellular pocket than the extracellular pocket, in agreement with the docking prediction that this compound preferentially interacts with the intracellular binding region. In contrast, Lidocaine was more strongly affected by mutation of the extracellular pocket, supporting a greater contribution of this region to receptor activation.

Tamoxifen showed a more complex profile. The initial S194A and S265A mutations produced relatively modest effects on receptor activation despite docking predictions suggesting interactions with both binding pockets. Additional mutations at F172 and F198, shifted the Tamoxifen concentration-response curves to higher concentrations while preserving substantial receptor activation. The limited effects of individual mutations, together with the retention of receptor activity across multiple mutant backgrounds, may indicate that Tamoxifen recognition relies on several receptor-ligand interactions rather than on a single dominant contact. However, additional structural studies will be required to determine the precise molecular basis of these effects.

Notably, only a limited set of residues in each binding pocket was probed in the present mutagenesis experiments, and additional positions may contribute to ligand recognition and signaling.

### 4.3 Physiological relevance of TAS2R14 activation by pharmaceutical agonists

We next considered whether the concentrations required for receptor activation are compatible with physiologically relevant exposure levels.

Exposure of TAS2R14 to pharmacologically relevant drug concentrations may occur both locally in the oral cavity and systemically following absorption. In addition, many drugs are known to partition from plasma into saliva, with reported saliva-to-plasma ratios typically ranging from 0.2 to 0.6, depending on physicochemical properties, indicating that circulating drugs may be continuously present in the oral cavity at measurable concentrations^36^.

Tamoxifen, administered orally, exhibits poor aqueous solubility (0.3 mg/L at pH 3–3.5), suggesting that only a fraction of an orally administered dose dissolves in saliva prior to swallowing; however, even limited dissolution could theoretically produce micromolar concentrations locally^37^. In plasma, Tamoxifen reaches steady-state concentrations of approximately 300 ng/mL (corresponding to 0.8 µM) after four weeks of daily treatment^38^, while its active metabolite endoxifen is present at lower plasma concentrations ranging from 16–173 nM^39^.

Carbimazole is rapidly converted to its active metabolite, methimazole, after absorption, indicating that Carbimazole itself is unlikely to be detected in plasma. Methimazole is reported to reach plasma concentrations of approximately 1.7–8.7 nM^40^ and has been experimentally shown not to induce TAS2R14 activation^41^. These observations suggest that any TAS2R14 activation associated with Carbimazole is more likely to occur before metabolic conversion, potentially during local exposure in the oral cavity.

In our experiments, Tamoxifen and Carbimazole were tested in concentration ranges of over 100 nM to 100 µM, whereas Lidocaine was tested at concentrations of 10 µM to 10 mM. Comparison of the measured EC_50_ values with reported drug exposure levels suggests that systemic TAS2R14 activation is likely limited for Tamoxifen and Carbimazole, whereas local exposure may be more relevant.

Lidocaine presents a different scenario, as local administration can achieve substantially higher tissue concentrations than those observed in circulation. The relatively high concentrations required for TAS2R14 activation are therefore more compatible with local rather than systemic exposure. Interestingly, TAS2R14-dependent apoptotic effects of Lidocaine have previously been reported in head and neck squamous cell carcinoma cells^17^, suggesting that TAS2R14-mediated effects are most likely to occur in tissues directly exposed to high local Lidocaine concentrations rather than through systemic drug exposure.

Notably, our estimates of physiological relevance should be viewed as order-of-magnitude indicators rather than precise predictions, and more detailed pharmacokinetic and tissue distribution data will be required to define exposure thresholds for TAS2R14 activation in vivo.

### 4.4 Conclusions

Pharmaceutical TAS2R14 agonists differ in potency, efficacy, signaling profiles and binding-site dependence. This highlights the importance of quantitative pharmacological characterization when evaluating clinically used drugs as TAS2R14 agonists.

By relating EC_50_ values to reported exposure levels, this study provides a basis for assessing when drug-induced TAS2R14 activation may be physiologically relevant. Such information may support future efforts to understand extraoral TAS2R14 functions and to evaluate existing drugs for potential TAS2R14-based repurposing.

## 5) Statements and Declarations

## Acknowledgements

We thank Professor Michael Naim for fruitful discussions.

## Author contributions

SE: Functional assays, writing and editing. ND: Functional and recruitment assays, writing and editing. AR and EZ: Computational analyses and visualization. EM: Supervision and project administration. MYN: Conceptualization, supervision, resources, writing and editing.

## Data Availability

The datasets generated during the current study are available from the corresponding author upon request.

