## Supplementary Figures for "Pharmaceutical TAS2R14 Agonists Display Diverse Potency, Efficacy, and Binding-Site Sensitivity"

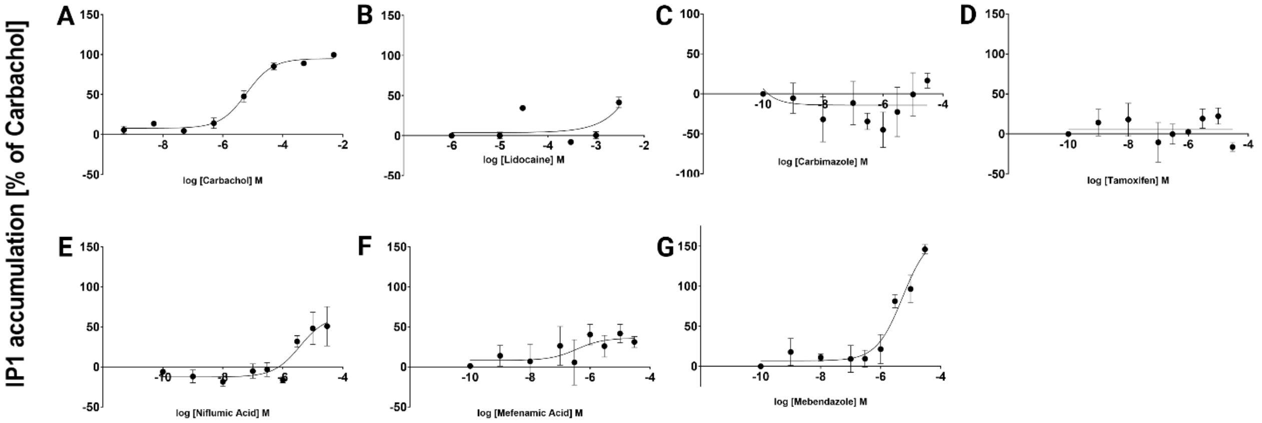


Figure S1: all tested compounds stimulation in HEK293 cells transfected with 2µg MOCK and 1µg Gαqi5 plasmids, measured Gαqi5 mediated accumulation of IP1% of Carbachol. Values represent the mean ±SEM. 3 biological repeats. (A) Carbachol- positive control (B) Lidocaine (C) Carbimazole (D) Tamoxifen (E) Niflumic acid (F) Mefenamic acid (G) Mebendazole.

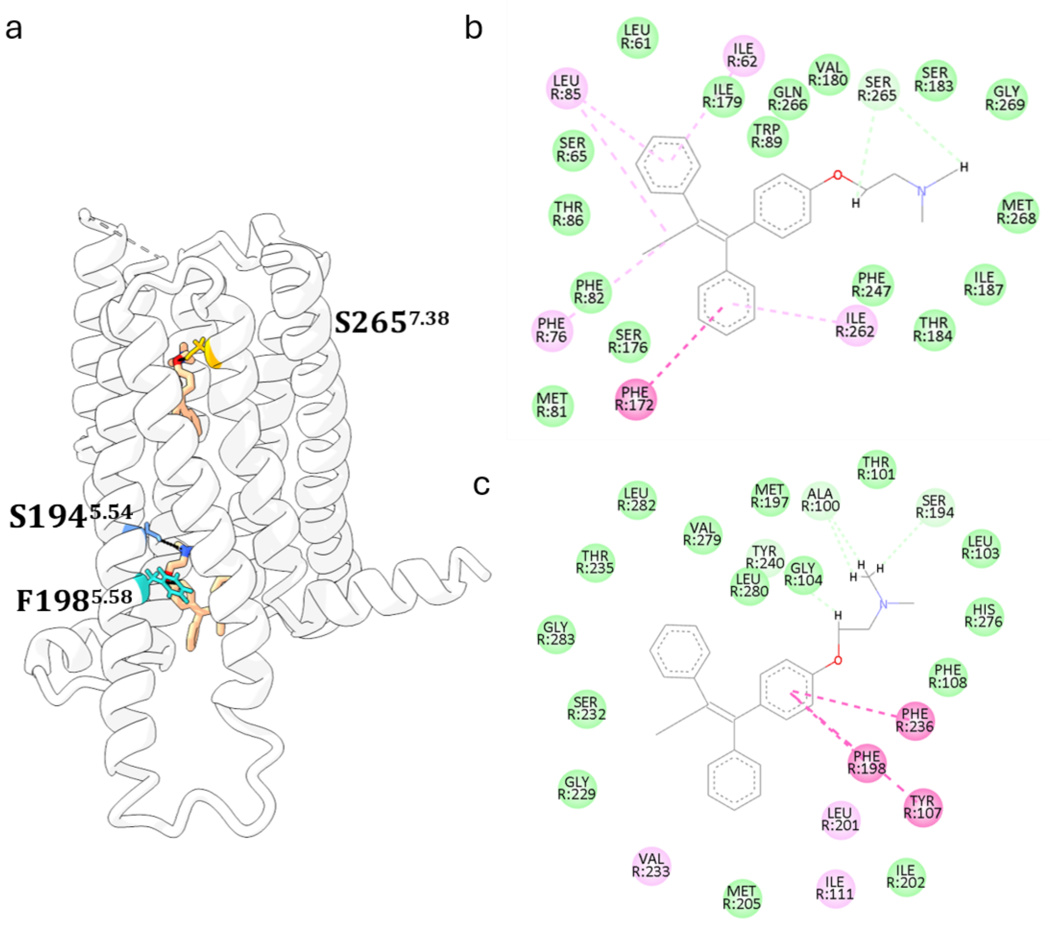


Figure S2: (a) TAS2R14 experimental structure (PDB: 8RQL) with the predicted docking pose of Tamoxifen. Serine 265 is shown in orange, Serine 194 in blue, and Phenylalanine 198 in turquoise. The figure was produced using ChimeraX. (b-c) Interaction maps of Tamoxifen in TAS2R14 extracellular (b) and intracellular (c) binding pockets. Protein-ligand interaction diagrams were generated using BIOVIA Discovery Studio Visualizer 2025.

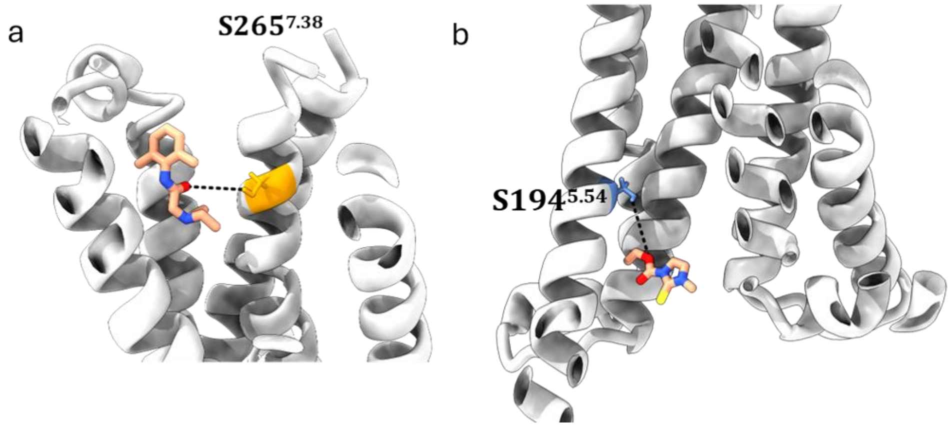


Figure S3: TAS2R14 experimental structure (PDB: 8RQL) with the predicted docking poses of Carbimazole and Lidocaine. (a) Predicted Lidocaine interactions within the extracellular binding pocket. (b) Predicted Carbimazole interactions within the intracellular binding pocket.

Serine 265 is shown in orange and Serine 194 in blue. The figure was produced using ChimeraX.
